# Deciphering Ca^2+^ Permeation and Valence Selectivity in Ca_V_1: Molecular Dynamics Simulations Reveal the Three-Ion Knock-on Mechanism

**DOI:** 10.1101/2024.11.27.625788

**Authors:** Lingfeng Xue, Nieng Yan, Chen Song

## Abstract

Voltage-gated calcium (Ca_V_) channels are pivotal in cellular signaling due to their selective calcium ion permeation upon membrane depolarization. While previous studies have established the highly selective permeability of Ca_V_ channels, the detailed molecular mechanism remains elusive. Here we use extensive atomistic molecular dynamics simulations to elucidate the mechanisms governing ion permeation and valence selectivity in Ca_V_1 channels. Employing the electronic continuum correction method, we simulated a calcium conductance of approximately 9–11 pS, aligning closely with experimental measurement. Our simulations uncovered a three-ion knock-on mechanism critical for efficient calcium ion permeation, necessitating the binding of at least two calcium ions within the selectivity filter (SF) and the subsequent entry of a third ion. *In silico* mutation simulations further validated the importance of multi-ion coordination in the SF for efficient ion permeation, identifying two critical residues, D706 and E1101, that are essential for the binding of two calcium ions in the SF. Moreover, we explored the competitive permeation of calcium and sodium ions, and obtained a valence selectivity favoring calcium over sodium at a ratio of approximately 35:1 under the bi-cation condition. This selectivity arises from the strong electrostatic interactions of calcium ions in the confined SF and the three-ion knock-on mechanism. Our findings provide novel insights into the molecular underpinnings of Ca_V_ channel function, with implications for understanding calcium-dependent cellular processes.

## 1 Introduction

Voltage-gated calcium (Ca_V_) channels mediate calcium influx upon membrane depolarization, initiating various cellular responses and playing a crucial role in many physiological functions such as muscle contraction and synaptic transmission. ^1,2^ Therefore, malfunctions of Ca_V_ channels can lead to a variety of severe diseases such as cardiac arrhythmias and ataxia, affecting both the nervous and cardiovascular systems. ^2^ Understanding the function mechanisms of Ca_V_ channels is crucial for developing therapeutic strategies to treat these conditions.

Ca_V_ channels are classified into three families: Ca_V_1, Ca_V_2, and Ca_V_3. Among these, Ca_V_1 channels, also known as L-type calcium channels, have been extensively studied for their high selectivity and permeability to Ca^2+^.^3^ The ion permeation characteristics of Ca_V_1 channels have been elucidated through electrophysiological experiments. Single-channel conductance measurements revealed that the Ca^2+^ conductance of Ca_V_1 is on the magnitude of 10 pS. ^4,5^ The anomalous mole-fraction behavior revealed two distinct affinities for Ca^2+^ in calcium channels: a micromolar affinity for the Ca^2+^ blockage of Na^+^ current and a millimolar affinity for saturation of Ca^2+^ current. ^6,7^ Reversal potential measurements showed that the permeability ratio of Ca^2+^ to monovalent ions can reach 1000:1. ^5,8–10^ However, in the absence of divalent ions, the conductance of monovalent ions can reach a conductance as high as 85 pS. ^5^ These observations led to the proposal of the two-ion pore model: ^6,7^ the channel can hold two Ca^2+^ ions, and efficient Ca^2+^ permeation occurs only when the channel is filled with two Ca^2+^ ions; when only one Ca^2+^ ion is present, the pore is blocked, preventing the permeation of monovalent ions.

Based on mutagenesis results, the key residues responsible for Ca^2+^ binding in Ca_V_1 were located at the EEEE locus in the selectivity filter (SF), ^11–13^ where four glutamate residues form a high-affinity site for calcium. No other high-affinity calcium-binding sites were identified. ^13,14^ Considering the two-ion pore model, these results suggested that the EEEE locus can offer two binding sites for calcium, probably resulting from the extended spacial distribution and the flexibility of side chains of the four glutamate residues. In addition to the EEEE locus, the divalent cation selectivity (DCS) locus was found as a low-affinity site, which influences divalent cation selection. ^15^

Although the general concept of the two-ion pore with a high-affinity EEEE locus has been established, the selectivity and permeation mechanisms of Ca_V_1 channels remain elusive. To address this issue, multiple theories and models have been proposed. Brownian dynamics simulations were used to investigate the calcium channel, discovering that the selectivity results from the strong electrostatic attraction of a divalent ion in the energy well, which prevents a monovalent ion from displacing it. ^16^ The charge/space competition mechanism proposed that the determinant of Ca^2+^ versus Na^+^ selectivity is the density of charged particles in the selectivity filter.^17^ The Coulomb blockade model explains the barrierless permeation of Ca^2+^ by the electrostatic balance of the bound ions and fixed charges at the SF. ^18^ These models significantly advanced the understanding of the ion permeation and selectivity mechanisms of Ca_V_ channels. However, a major limitation still exists in that those models were not based on the actual high-resolution channel structure, hindering an accurate and quantitative description of the ion permeation process.

Recently, the atomistic structures of Ca_V_1 channels have been resolved by cryo-EM, ^19–21^ which provide the structural basis for understanding Ca_V_1 functions. With these structures, molecular dynamics (MD) simulations can be employed to investigate the ion binding, permeation, and selectivity mechanisms of Ca_V_1 channels. ^22,23^ Using MD simulations, Zhu et al. identified two Ca^2+^ binding sites at the EEEE locus, ^23^ in line with previous studies. However, further MD investigations on the ion permeation and selectivity of Ca_V_ channels have been hindered by the inability of the classical force fields to account for charge transfer and polarization, which therefore often overestimates the binding affinities of divalent ions to proteins. ^24^ This is also one of the reasons that extensive MD simulations have been conducted for voltage-gated potassium and sodium channels, ^25–29^ but rarely for calcium channels. ^30–33^

The limitations of the classical force fields have been frequently observed in MD simulations of ion channels, which require highly accurate descriptions of the interactions between ions and proteins. For instance, in the gramicidin A channel, the energy barrier for ion permeation was overestimated, yielding a conductance value five orders of magnitude less than the experimental value. ^34^ Similarly, in potassium channels, the channel conductance under physiological membrane potential was found to be too low ^35^ and the energy barrier for ion permeation too high. ^36^ To address these limitations, polarizable force fields ^37^ have been developed and tested. Two polarizable force fields, AMOEBA and Drude, have been applied to the gramicidin A channel, ^38,39^ both yielding more reasonable energy barriers than the non-polarizable force fields. However, the polarizable force fields are less efficient and often require more computational power and time. Therefore, researchers have also been trying to modify the non-polarizable force fields to improve the model’s accuracy. For example, we utilized the multi-site strategy and developed a new Ca^2+^ model, ^33^ which was able to quantitatively simulate Ca^2+^ permeation and selectivity in ryanodine receptors. ^33,40^ However, in the narrower TRPV channels, the model still needs a high voltage to generate a reasonable Ca^2+^ conductance. ^41,42^ Another promising strategy is the electronic continuum correction (ECC) method developed by Leontyev et al. ^43^ Recently, both polarizable force field and ECC method were applied to study the ion occupancy in the KcsA channel, consistently supporting a thermodynamically stable four-ion configuration. ^44^ Also, the ECC method has been applied to another potassium channel MthK, giving a reasonable conductance. ^45^

In this study, we used the ECC method to correct the CHARMM force field and obtained a series of simulation results that aligned well with existing experimental data, including the Ca^2+^ conductance and the residue mutation effects. More importantly, our simulations revealed the three-ion knock-on mechanism for Ca^2+^ permeation in Ca_V_1 channels and identified the key residues responsible for the fast Ca^2+^ permeation. Moreover, we observed the selectivity of Ca^2+^ over Na^+^ at a ratio of about 35 in the MD simulations. Our results indicate that the valence selectivity of Ca_V_1 channels can be attributed to the strong electrostatic interactions of Ca^2+^ at the confined SF region formed by negatively charged residues, as well as the three-ion knock-on mechanism.

## 2 Results

### 2.1 Structural model and simulation protocol validation

Although several Cryo-EM structures of Ca_V_ channels have been determined, ^19–21^ an open-state structure is not available yet. Therefore, to investigate the ion permeation mechanism in Ca_V_1, we computationally built an open-state structure based on the closed-state Cryo-EM structure of Ca_V_1.3, ^21^ with the gate region modeled according to an open-state Na_V_ structure. ^46^ Specifically, the S6 helices of the open-state Na_V_ structure were aligned to Ca_V_1.3 and used as the reference structure for position restraints in the MD simulations. By adding restraints to the intracellular gate region of the S6 helices, the hydrophobic gate of the Ca_V_1.3 structure became dilated, hydrated, and permeable for ions (Fig. S1). Notably, the conserved asparagine residues (residue numbers 399, 745, 1145, and 1457 in human Ca_V_1.3) were reoriented to face the pore, suggesting an open state similar to Na_V_ channels. ^46^ Such modeling was based on the assumption of structural similarity between the open-state Ca_V_ and Na_V_ channels, as well as the assumption that the SF structure will not be significantly changed by the dilation of the intracellular gate.

Using the open-state model, we performed ion permeation simulations with the default CHARMM force field and observed that Ca^2+^ got stuck in the SF, a phenomenon consistent with previous MD simulations for Ca_V_1 channels. ^23^ This was attributed to the inherent limitations of the classical nonpolarizable force field in accurately representing Ca^2+^-protein interactions. ^24^ Given that even the recently developed multisite Ca^2+^ model tended to overestimate the binding affinity of Ca^2+^ in narrow pores, ^41^ we opted to employ the ECC method, which effectively modulates the electrostatic interactions by applying a scaling factor between charged groups during simulations. The scaling factor for ECC was calibrated by fitting the experimental conductance of Ca_V_1. Through a series of ion permeation simulations conducted at a voltage of approximately 100 mV (Movie S1), we observed a conductance reduction with an increasing scaling factor (Fig. 1A). Notably, a scaling factor of 0.87 was identified as optimal, yielding a channel conductance of 9 pS at 300 K (Fig. 1A). This value is in good agreement with the experimentally determined range of 9 to 10 pS. ^5,47^ Therefore, this scaling factor was selected for the subsequent further validation and in-depth investigation in our MD simulations.

**Figure 1:**
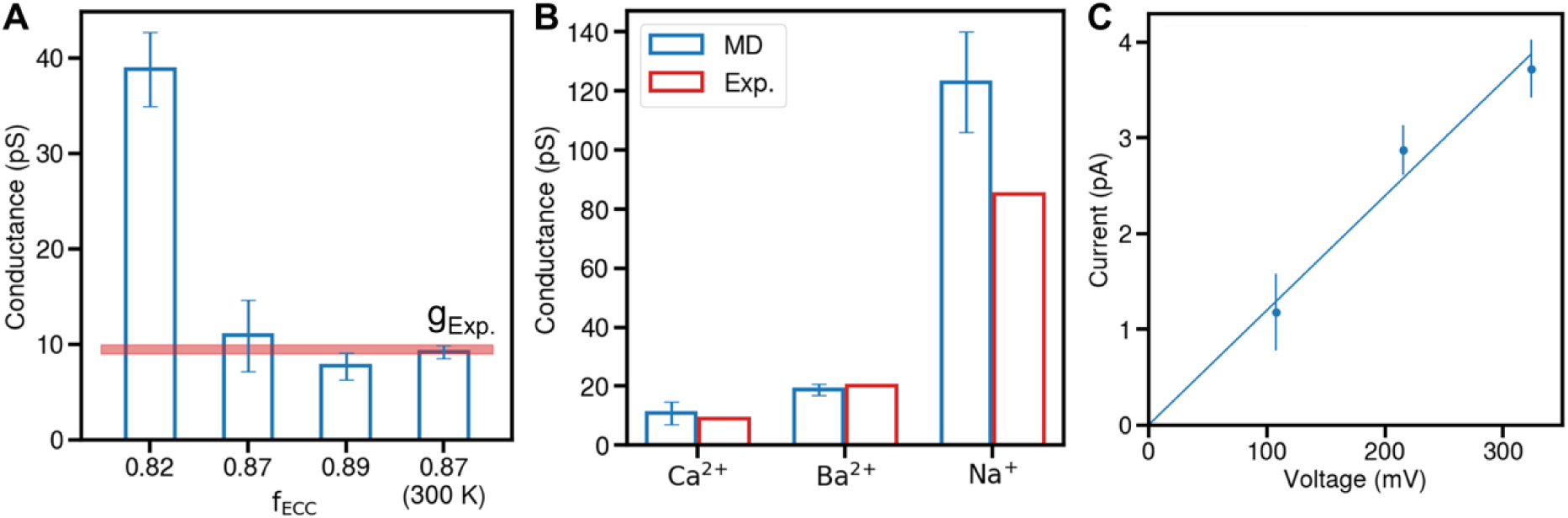
Conductance validation for the simulation system and protocol. (A) Ca^2+^ conductance of Ca_V_1 in our MD simulations with varying scaling factors for the ECC method at 310 K, with an additional simulation performed at 300 K. The red line represents the experimental conductance range for reference. (B) Simulated conductance values for three types of cations (blue) compared to experimental data (red). All simulations of (A) and (B) were conducted at a transmembrane voltage of approximately 100 mV. (C) The current-voltage relationship for Ca^2+^. The blue line represents a linear regression fit through the origin, indicating the channel’s ohmic behavior in the MD simulations.

With the scaling factor of 0.87, we expanded the ion permeation simulations to Ba^2+^ and Na^+^, and found that the Ba^2+^ conductance is about 2-fold of Ca^2+^, while the Na^+^ conductance is approximately 11-fold of Ca^2+^ at a voltage of 100 mV (Fig. 1B). The overall conductance sequence is Na^+^ ≫ Ba^2+^ > Ca^2+^, which is consistent with experimental measurements, ^5^ although the absolute value of Na^+^ conductance is slightly overestimated in our simulations. One concern regarding the conductance in MD simulations is the potential nonlinearity of the current-voltage relationship, ^35^ which often makes the conductance incompatible with experimental measurements. To address this, we explored the I-V curves for the three cations. Reassuringly, the I-V relations are linear for all three cations within the physiological voltage range (Fig. 1C, Fig. S2A), although the I-V curve for Ba^2+^ becomes superlinear at higher voltages (Fig. S2A). These results support that our structural model, simulation setup, and simulation protocol can reproduce the major physiological characteristics of Ca_V_1.

In addition to assessing ion conductance, we conducted a comparative analysis of ion density within the SF of Ca_V_1.3, leveraging both our computational results and the closed-state cryo-EM density. For this purpose, we used the closed structure of Ca_V_1.3 to conduct a 500-ns simulation with one Ca^2+^ in the SF. With the EEEE locus center as the reference origin, the primary binding site for Ca^2+^ along the z-axis (membrane normal) was located between -1 to 0 Å, in close proximity to the experimental density peak observed at around 0 Å (Fig. S2B, C), further validating the accuracy of the ECC scaling factor. To identify the binding sites when two Ca^2+^ ions are present in the SF, we performed two-Ca^2+^ Metadynamics simulations. The results revealed two predominant binding sites located at -2.8 Å and 1.7 Å along the z-axis (Fig. S2D). Notably, these binding sites differ from the binding site observed in the presence of a single Ca^2+^ ion, indicating a different binding pattern when two Ca^2+^ ions are concurrently present within the SF.

### 2.2 Ca^2+^ occupancy states in the SF

With the validated simulation system, we performed 37.5-μs ion permeation simulations under approximately 100 mV to investigate the Ca^2+^ permeation mechanism in Ca_V_1 (Movie S2). A detailed analysis of the Ca^2+^ density distribution around the SF revealed five binding sites for Ca^2+^ (Fig. 2A, B): S1U, S1L, S2U, and S2L within the SF (where S denotes the selectivity filter, U denotes upper, and L denotes lower), along with V1 in the outer vestibule (V denotes vestibule). Importantly, due to the mutual exclusivity of binding within the S1U/S1L and S2U/S2L pairs, our simulations indicate that a maximum of three Ca^2+^ ions can engage these five sites most of the time. Specifically, we observe two Ca^2+^ ions bound simultaneously within the SF, with a third ion located in the outer vestibule above the SF.

**Figure 2:**
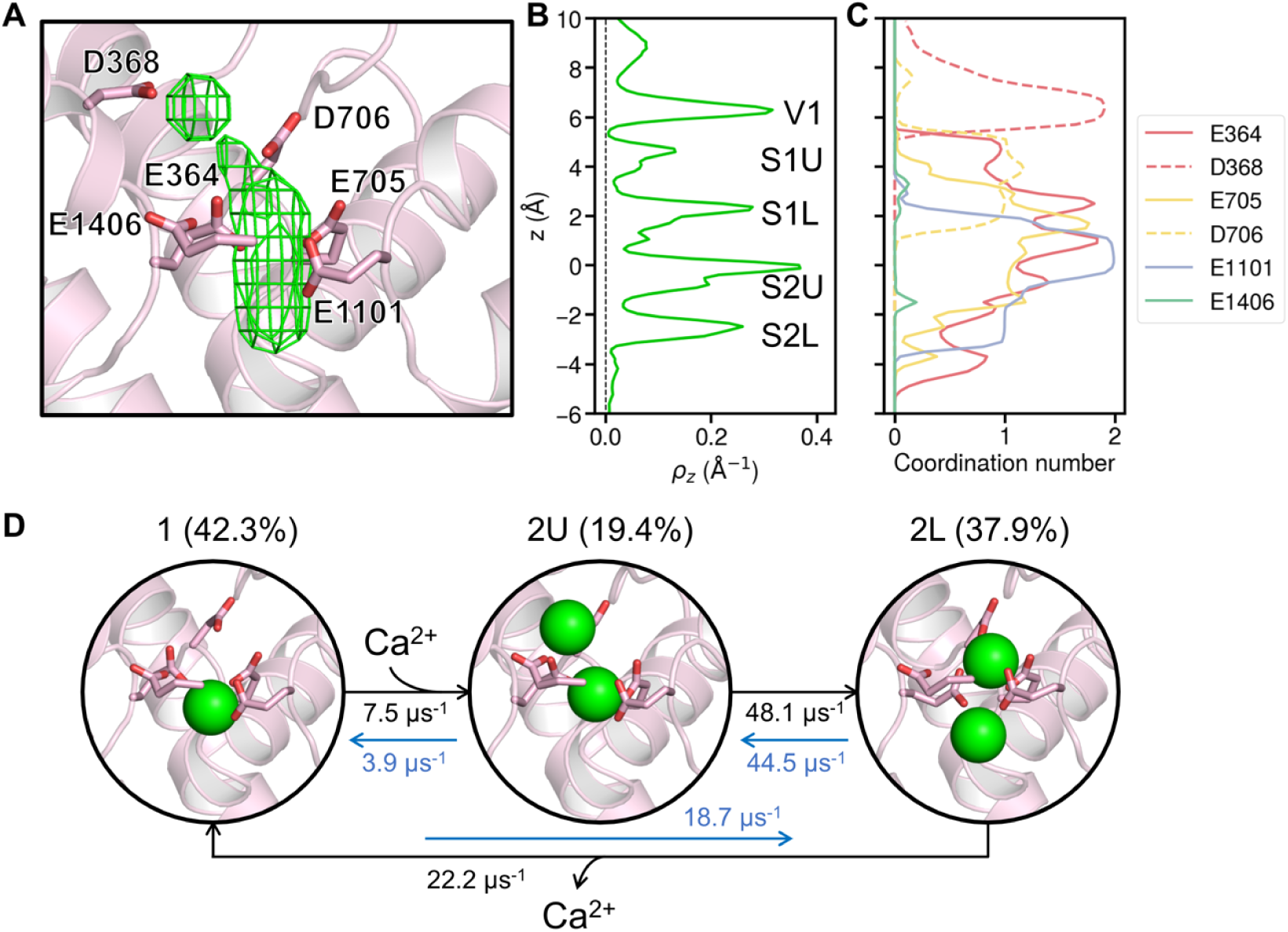
Ca^2+^ binding and coordination within the selectivity filter (SF) of Ca_V_1. (A) Density map of Ca^2+^ ions (green grids) within the SF. The protein backbone is shown as cartoon, with repeat IV omitted for clarity. The side chains of key residues are shown as sticks, with residue numbers labeled. (B) Z-distribution of Ca^2+^ ions from ion permeation simulations, with the binding sites annotated corresponding to the major peaks. (C) Coordination number distribution of Ca^2+^ ions with the carboxylate oxygens of the key residues in the SF along z-axis. Residues from the four repeats are colored red, yellow, blue, and green, respectively. (D) Schematic diagram illustrating the three major Ca^2+^ binding patterns and the permeation cycle within the SF, with the occupancy probabilities and transition rates labelled for the three states. Green spheres represent Ca^2+^ ions.

The binding sites within the SF are orchestrated by the electrostatic interactions with the surrounding acidic residues. Our analysis of the coordination patterns of Ca^2+^ ions as they permeate the SF revealed that each ion typically sheds up to five coordinated water molecules, forming direct contact with the carboxylate oxygens of the SF residues (Fig. S3A). In contrast, coordination with carbonyl oxygens was minimal (Fig. S3A). A detailed residue-level coordination analysis pinpointed the key contributors to each binding site (Fig. 2C). Specifically, at the V1 site in the outer vestibule, the Ca^2+^ ion exclusively coordinates with the aspartate residue D368, establishing the DCS locus. In the S1U and S1L sites, coordination occurs with two glutamate residues E364 and E705 and the aspartate residue D706, underscoring the significant role of D706 in Ca^2+^ binding. Aligning with recent literature, ^23,48,49^ our findings confirm that the D706 residue, in addition to the EEEE locus, is integral to Ca^2+^ binding within the SF, effectively extending the binding motif to an EEEED locus. At the S2U and S2L sites, the Ca^2+^ ion exhibits an asymmetric coordination pattern, engaging with E364, E705 and E1101 of the EEEE locus.

Notably, the coordination of Ca^2+^ by the four glutamate residues within the EEEE locus exhibits significant asymmetry. Our simulations revealed that the E364 residue, owing to the flexibility of its side chain, engages in a broad range of coordination interactions, whereas E705 and E1101 demonstrate a more focused coordination profile (Fig. 2C). Notably, E1101 exhibits bidentate coordination with Ca^2+^ around S2U site, suggesting its key contribution to Ca^2+^ binding stability. In contrast, the E1406 residue was found to rarely directly coordinate with Ca^2+^ ions (Fig. 2C, green line). The distinct conformation of the E1406 side chain, which diverges from the other glutamate residues, appears to be a key factor for the Ca^2+^ coordination difference. Unlike the other residues that are oriented towards the pore axis, E1406 is positioned towards the backbone of E364, forming hydrogen bonds with the backbone nitrogen atoms (Fig. S3B). This unique orientation suggests that the fourth glutamate of the EEEE locus may play a diminished role in direct Ca^2+^ coordination.

During the ion permeation process, the four Ca^2+^ binding sites in the SF form three distinct ion binding patterns, which alternate to form a continuous ion permeation cycle. By examining the coordination of Ca^2+^ ions with the EEEED locus and their z-axis distributions, we identified three major states for Ca^2+^ permeation: state 1, 2U, and 2L (Fig. 2D). In state 1, a single Ca^2+^ ion occupies the SF, predominantly at the S2U site, where it coordinates with the first three glutamates of the EEEE locus. State 2U is characterized by two Ca^2+^ ions binding to the upper sites, S1U and S2U, whereas in state 2L, the ions bind to the lower sites, S1L and S2L. The permeation cycle can then be described as follows (Fig. 2D):

1. The cycle initiates with a single Ca^2+^ ion situated in the SF, coordinating with three glutamates at the S2U site.
2. The approach of an additional Ca^2+^ ion from the outer vestibule instigates a transition; the ion at the S2U site remains in place, while the new ion settles into the S1U site.
3. Subsequent cooperative translocation results in both ions shifting to the lower sites, S1L and S2L.
4. Ultimately, the lower Ca^2+^ ion exits the SF into the central cavity, reinstating the SF with a single Ca^2+^ ion and reverting to the original state 1.

By analyzing the occupancy probabilities of the three states, we found that states 1 and 2L exhibit higher probabilities than state 2U in these ion permeation simulations. Further analysis of transition rates showed that Ca^2+^ entry is the most irreversible step: once that the Ca^2+^ ion enters the SF from outer vestibule, it is unlikely to move back (Fig. 2D).

### 2.3 Ca^2+^ permeation mechanism

To elucidate the Ca^2+^ permeation mechanism in Ca_V_1, we employed Metadynamics simulations to quantify the energetics of the permeation process, thereby explaining the high conductance of Ca^2+^ in Ca_V_ channels. The collective variables (CVs) utilized in our analysis were the z-positions of Ca^2+^ ions relative to the EEEE locus. We conducted potential of mean force (PMF) calculations for systems containing one, two, and three Ca^2+^ ions.

The PMF results for the single-Ca^2+^ system exposed a substantial energy well within the SF (Fig. S4A), with an exit energy barrier of approximately 39 kJ/mol for Ca^2+^, a value too high to facilitate ion permeation. This implies that permeation with a single Ca^2+^ ion in the SF is improbable, aligning with the established two-ion pore model. ^6,7^

In the two-Ca^2+^ PMF, the energy barrier for Ca^2+^ exit was calculated to be approximately 27 kJ/mol (Fig. S4B-D), which, while lower, remains a significant barrier to efficient permeation. Examination of ion configurations along the minimum free energy path (MFEP) indicated that this elevated energy barrier materializes at a reaction coordinate of around 0.8 (Fig. S4C-D). This configuration corresponds to an unfavorable Ca^2+^ ion occupancy state 1 (Fig. 2D), specifically when the lower Ca^2+^ ion departs from the SF, leaving the upper ion at the S2U site (z ∼ 0 Å) unable to neutralize the substantial negative charges of the SF carboxylates. This observation suggests that state 1 is not energetically favorable in a two-ion system. Therefore, the inclusion of additional ions in PMF calculations is necessary to fully unravel the mechanism of ion permeation.

In Metadynamics simulations under the three-Ca^2+^ condition, the MFEP revealed a substantially reduced energy barrier of approximately 14 kJ/mol (Fig. S4E, Fig. 3A) for Ca^2+^ ion exit. This finding is consistent with the rapid Ca^2+^ permeation characteristic of Ca_V_ channels. Analysis of ion configurations along the MFEP demonstrated that states 2U and 2L are energetically favorable, whereas state 1 exhibits a higher free energy of approximately 10 kJ/mol (Fig. 3A). The energy barriers between these three states are also reasonably minimal, facilitating rapid state transitions. In contrast to the two-Ca^2+^ PMF (Fig. S4B-D), state 1 in the three-Ca^2+^ PMF exhibits a significantly lower energy profile. This reduction occurs because, upon the exit of the lower Ca^2+^ ion from the SF, an additional ion from the outer vestibule provides stabilization through electrostatic interactions (Fig. 3C, state 1). Thus, our findings indicate that the presence of one or two Ca^2+^ ions is inadequate to account for efficient ion permeation in Ca_V_ channels. Instead, the inclusion of a third Ca^2+^ ion is essential, corroborating the three-ion knock-on mechanism as pivotal for understanding permeation efficiency.

**Figure 3:**
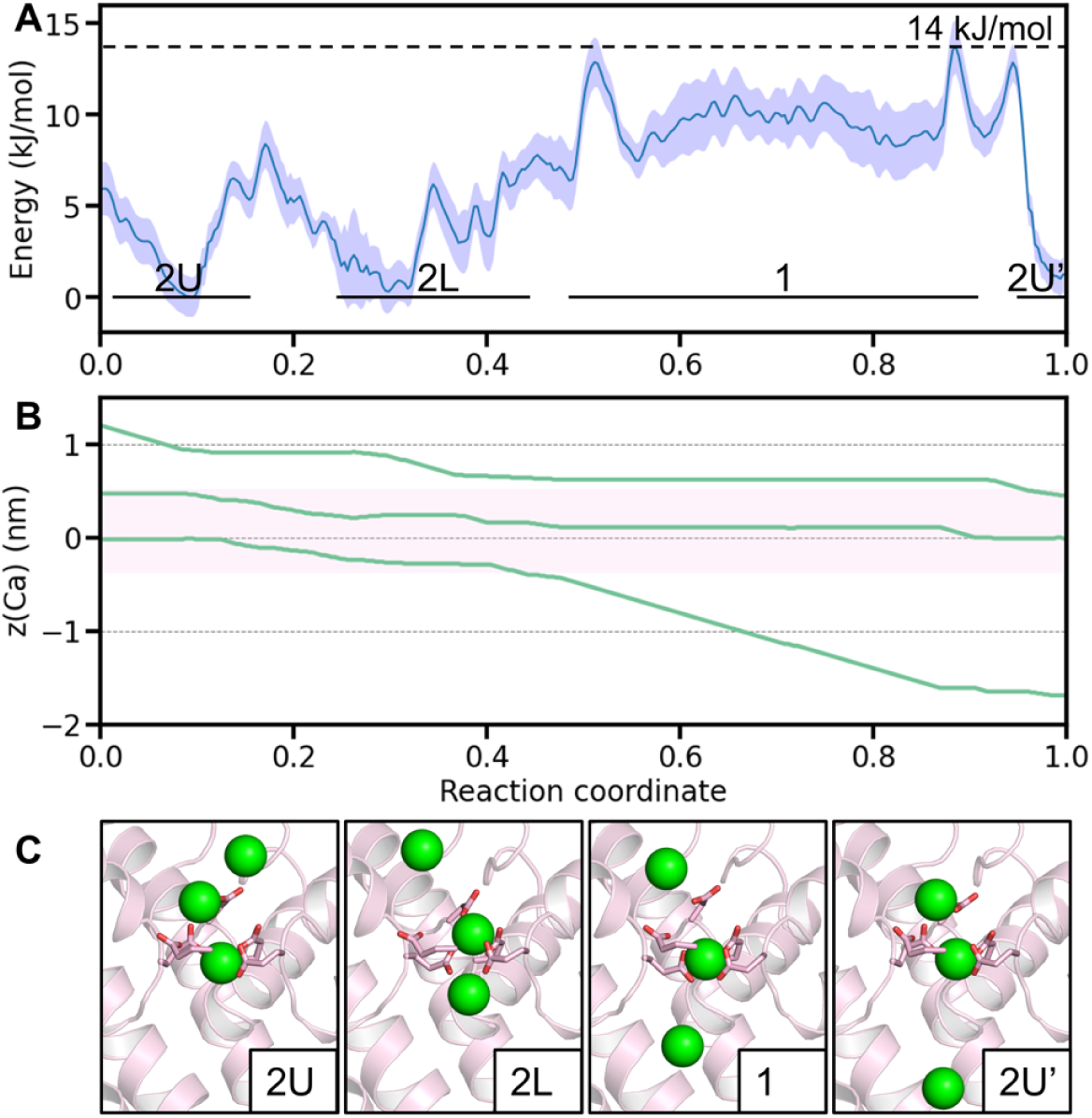
Permeation mechanism of Ca_V_1 revealed by three-Ca^2+^ PMF calculations. (A) Free energy profile along the MFEP of the three-ion PMF, with shaded area representing error bars. The different states are marked in the panel according to their corresponding z-positions in (B). (B) Coordinated motion of three Ca^2+^ ions along the z-axis of the MFEP. The pink shaded area indicates the SF region. (C) Representative SF configurations for the four key states during Ca^2+^ permeation. Side chains of the EEEED residues are shown as sticks. Green spheres represent Ca^2+^ ions.

It is interesting to note that the relative occupancy, i.e., the relative stability of the three states are different in the ion permeation simulations and three-ion PMF calculations (Fig. 2D, Fig. 3A). This is probably due to the fact that the former is non-equilibrium simulations while the latter is equilibrium simulations, which will be further discussed below.

To pinpoint the key residues facilitating rapid ion permeation in Ca_V_1, we conducted ion permeation simulations on five single-site mutants, each with a critical acidic residue at the EEEED locus mutated to alanine. Alongside Ca^2+^, we expanded our investigation to include Ba^2+^ simulations for a comparative analysis with existing experimental data. ^49–51^

Our simulations identified D706 and E1101 as the primary determinants of divalent cation conductance in Ca_V_1. Mutations at these positions substantially diminished conductance (Fig. 4A, Table 1). These residues, D706 at the top and E1101 at the bottom of the SF, are critically positioned to generate an extensive region of negative electrostatic potential, essential for stabilizing the two-Ca^2+^ configurations within the SF. Mutations at these sites result in a constriction of the Ca^2+^ binding region, disrupting the two-ion configuration and consequently reducing conductance. This hypothesis is corroborated by ion density analysis (Fig. 4C), which shows a reduction to a single binding site for Ca^2+^ in the D706A and E1101A mutants, thus precluding the two-ion configuration. In contrast, other mutants maintain at least two binding sites, preserving fast ion permeation. Again, this supports the three-ion knock-on permeation mechanism.

**Figure 4:**
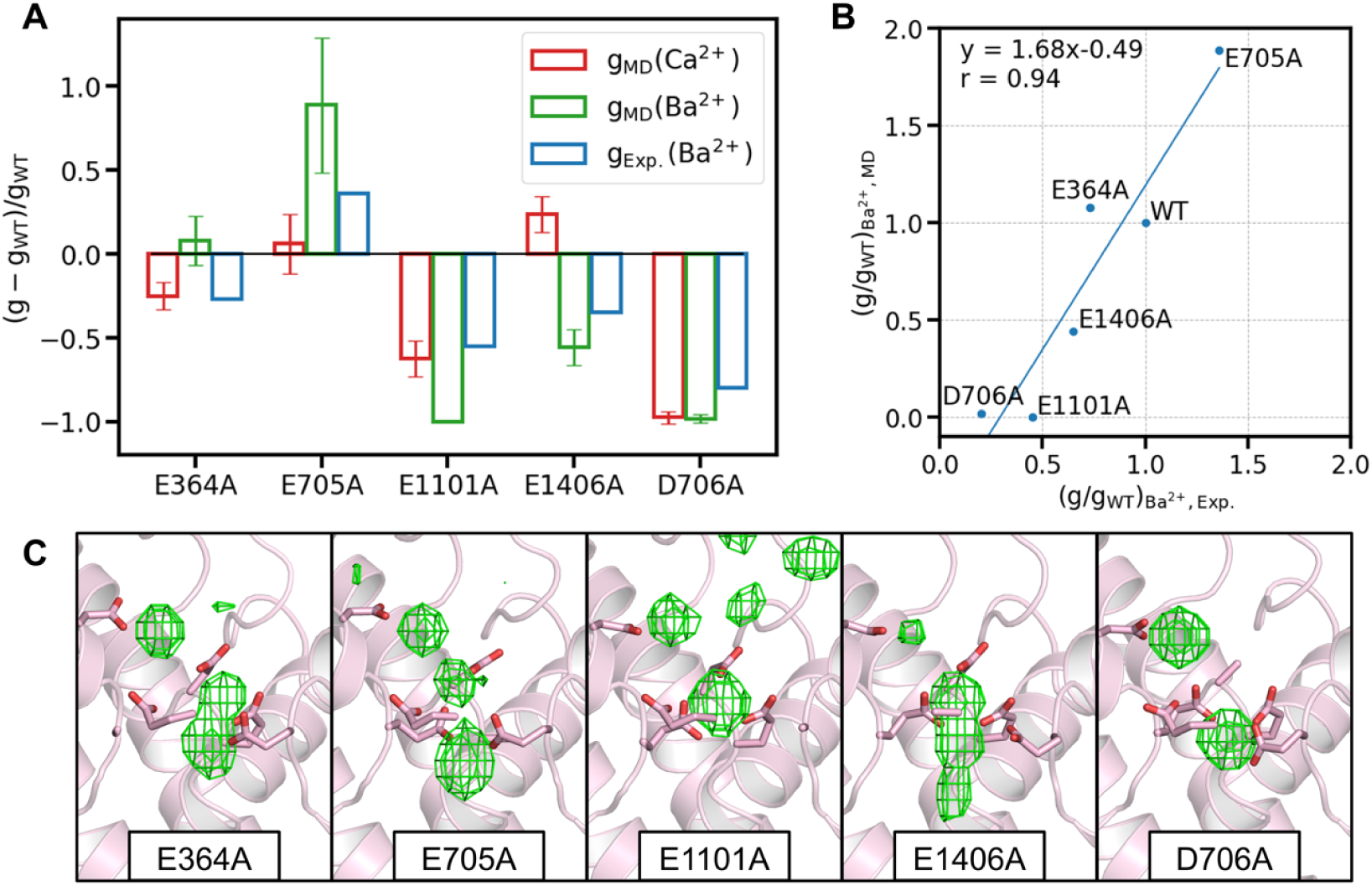
Conductance and Ca^2+^ density for Ca_V_1 mutants. (A) Relative conductance change for the five mutants, for Ca^2+^ (red) and Ba^2+^ (green) in simulations. Experimental data for Ba^2+^ (blue), from literature, ^49,50^ are also shown for comparison. (B) Ba^2+^ conductance fold-change for the mutants, plotted against experimental data. The blue line represents linear fitting results, with the fitted equation and Pearson’s coefficient displayed. (C) The Ca^2+^ density map (green grids, isosurface value=0.12) in the SF for the five mutants. Side chains of key residues are shown as sticks.

**Table 1:** Mutant conductance results.

| Mutant | $(g/g_{WT})_{MD, Ca}$ | $(g/g_{WT})_{MD, Ba}$ | $(g/g_{WT})_{Exp., Ba}$ |
| --- | --- | --- | --- |
| E364A | $0.75 \pm 0.08$ | $1.08 \pm 0.15$ | $0.73^{50}$ |
| E705A | $1.06 \pm 0.18$ | $1.89 \pm 0.40$ | $1.36,^{50} 1.49^{51}$ |
| E1101A | $0.37 \pm 0.11$ | $0 \pm 0$ | $0.45^{50}$ |
| E1406A | $1.23 \pm 0.11$ | $0.44 \pm 0.11$ | $0.65,^{50} 0.84^{51}$ |
| D706A | $0.03 \pm 0.04$ | $0.02 \pm 0.02$ | $0.2^{49}$ |

Comparison of our simulation results with experimental data revealed a strong concordance in the trend of Ba^2+^ conductance changes, with a Pearson’s correlation coefficient of 0.94 (Fig. 4B), although the simulations tended to overestimate the absolute conductance changes in mutants. For Ba^2+^, the general trend of mutation effects mirrored that of Ca^2+^, with the notable exception of the E1406A mutant, which unexpectedly increased Ca^2+^ conductance while decreasing Ba^2+^ conductance. While a definitive explanation is yet to be determined, we hypothesize that this anomaly may arise from the size discrepancy between the hydrated Ba^2+^ and Ca^2+^.

### 2.4 Ca^2+^ selectivity mechanism

The pronounced selectivity for Ca^2+^ over monovalent cations is a hallmark characteristic of Ca_V_ channels. To quantitatively assess this selectivity, we performed both bi-cation and mono-cation simulations. The bi-cation simulations included the presence of both Ca^2+^ and Na^+^ ions, whereas the mono-cation simulations involved either Na^+^ or Ca^2+^ ions exclusively. Our findings revealed that the Ca^2+^ conductance was not influenced by the coexistence of Na^+^ ions. In contrast, the Na^+^ conductance was markedly reduced to approximately 1/800 of its mono-cation value in the presence of Ca^2+^ (Fig. 5A). This substantial decrease in Na^+^ conductance is attributed to the obstructive presence and blocking effect of Ca^2+^ ions, as evidenced by the SF configurations and Na^+^ density analysis under bi-cation condition (Fig. 5B, C). Under mono-cation Na^+^ condition, the SF could accommodate 3–4 Na^+^ ions, enabling rapid Na^+^ permeation (Movie S3). However, in the presence of Ca^2+^, the SF was predominantly occupied by Ca^2+^ ions, resulting in a significantly diminished Na^+^ density (Fig. 5C). In our bi-cation simulations spanning 37.5 microseconds, we recorded 140 Ca^2+^ permeation events versus only 4 Na^+^ events (Movies S4-6), yielding a permeation event ratio of 35:1. This ratio reproduces the pronounced preference of Ca_V_1 for Ca^2+^ over Na^+^, which is captured in MD simulations for the first time.

**Figure 5:**
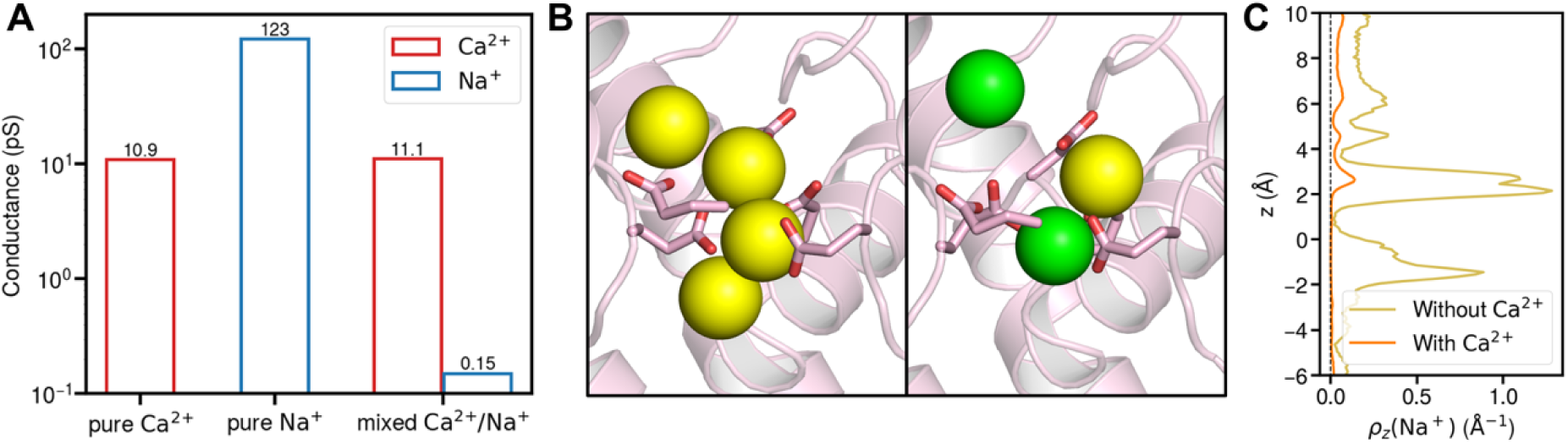
Ion permeation simulations for Ca_V_1 under mono-cation and bi-cation conditions. (A) Conductances of Ca^2+^ (red) and Na^+^ (blue) under three ionic conditions: pure Ca^2+^, pure Na^+^, and mixed Ca^2+^/Na^+^. (B) The typical configurations of Na^+^ in the selectivity filter under conditions of pure Na^+^ (left) and mixed Ca^2+^/Na^+^ (right). Side chains of the EEEED residues are shown as sticks. Green spheres represent Ca^2+^ ions and yellow spheres represent Na^+^ ions. (C) The z-distributions of Na^+^ in the absence (brown) and in the presence of Ca^2+^ (orange).

To understand the selectivity mechanism of Ca_V_1, we calculated the one-Na^+^-two-Ca^2+^ PMF, in which two Ca^2+^ ions were put in the SF while a third Na^+^ ion was trying to push through (Fig. 6). From the resulting three-dimensional PMF (Fig. S4F), we constructed the MFEP and found that the energy barrier for Na^+^ permeation through SF is ∼33 kJ/mol (Fig. 6A), approximately 19 kJ/mol higher than that of Ca^2+^ permeation. According to the Arrhenius law, this energy difference corresponds to a rate ratio of ∼1600:1, an extremely high selectivity for Ca^2+^. It should also be noted that the MFEP does not represent a three-ion knock-on process. Rather, it appears that the permeating Na^+^ bypasses the two bound Ca^2+^ ions to translocate (Fig. 6C).

**Figure 6:**
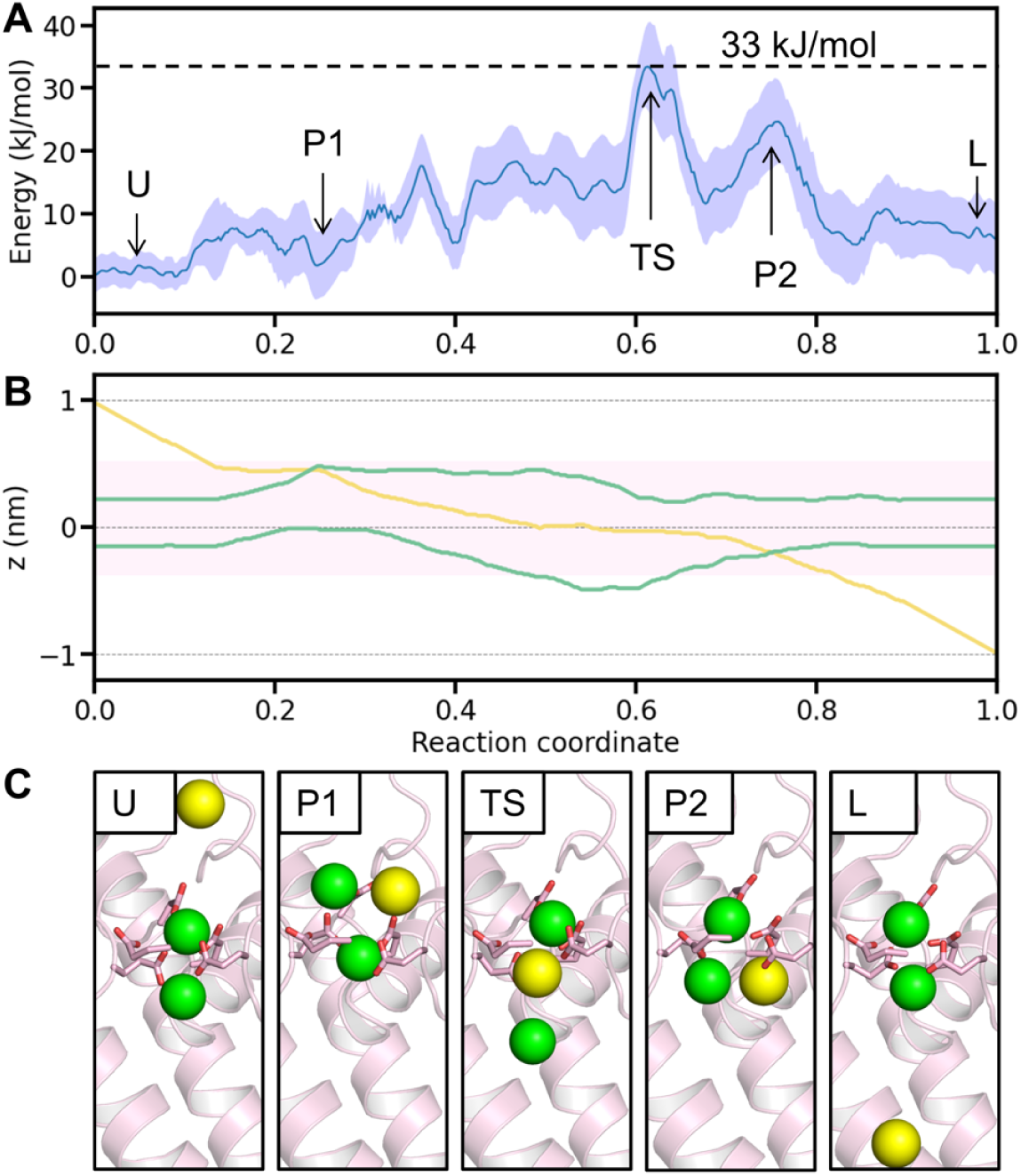
Unfavorable Na^+^ ion permeation revealed by one-Na^+^-two-Ca^2+^ PMF calculations. (A) Free energy profile along the MFEP, with shaded area representing error bars. The different states are marked according to the corresponding z-positions in (B). (B) Motion of one Na^+^ (yellow) and two Ca^2+^ ions (green) along the z-axis of the MFEP. The pink shaded area indicates the SF region. (C) Representative SF configurations for the five key states during Na^+^ permeation. Side chains of the EEEED residues are shown as sticks. Green spheres represent Ca^2+^ ions and yellow spheres represent Na^+^ ions.

In the one-Na^+^-two-Ca^2+^ PMF, the most stable configuration consists of two Ca^2+^ ions bound in the SF with a Na^+^ ion outside (Fig. 6C, states U and L). When Na^+^ enters the SF and passes the upper Ca^2+^ ion (state P1), the energy remains manageable due to the Na^+^ ion coordinating with D706 and E1101. However, as Na^+^ moves downward, pushing the lower Ca^2+^ ion out of the SF, the energy spikes due to the unfavorable electrostatic interactions upon the lower Ca^2+^ moving away from the SF, corresponding to the transition state for Na^+^ permeation (state TS). Upon passing the lower Ca^2+^ ion (state P2), the energy remains unfavorable, in contrast to state P1. This is probably due to the fact that not enough negatively charged residues can balance the coexistence of the Ca^2+^ and Na^+^ ions below the SF. After Na^+^ exits, the lower Ca^2+^ ion returns to its initial site. As can be seen, the MFEP represents a bypass permeation mechanism for Na^+^ in the presence of two bound Ca^2+^ in the SF, and the free energy barrier is too high to allow rapid permeation.

Note that compared with the three-Ca^2+^ PMF (Fig. S4E), many more paths were found in one-Na^+^-two-Ca^2+^ PMF (Fig. S4F), indicating that there may be many different pathways with similar energy barriers for Na^+^ permeation. However, the pathways around the transition state are converged (Fig. S4F), indicating that the exit of the Na^+^ from the SF is always the limiting step (Fig. 6C, state TS). The high energy of the transition state arises from the unfavorable electrostatic interactions between the three stacked ions and the SF. Specifically, before reaching the transition state, the permeating Na^+^ and the upper Ca^2+^ cannot provide enough repulsion to push away the lower Ca^2+^, which is still attracted to the SF. Consequently, the Na^+^ has to bypass the lower Ca^2+^ to permeate rather than pushing and following the downward motion of the lower Ca^2+^ to form a knock-on permeation. This is evident from the Fig. 6B where the yellow line crossing the lower green line, corresponding to state P2 in Fig. 6C. Although such a bypass permeation is more favorable for the one-Na^+^-two-Ca^2+^ system, its energy barrier is much higher than that of the three-Ca^2+^ knock-on permeation, which can yield a selectivity of Ca^2+^ over Na^+^ permeation.

When comparing the selectivity ratios from ion permeation simulations (Fig. 5A) and PMF calculations (Fig. 3A, Fig. 6A), we observed significant discrepancies. The former indicated a permeation event ratio of 35:1, while the latter was approximately 2000:1 according to the Arrhenius law. This discrepancy may represent the differences and limitations of the two methods. First of all, the former is non-equilibrium simulations, while the latter a very rough estimation from equilibrium simulations. The former is an observation from simulations that took into account the effects of all the surrounding ions and possible permeation pathways, while the latter only considered three ions and one particular permeation pathway (MFEP). In fact, we did observe concerted translocation of two Na^+^ ions through the SF in the simulated bi-cation permeation trajectories (Movie S6), highlighting the limitation of considering three-ion MFEP only. In addition, the application of a high voltage (100 mV) likely reduced the selectivity in the ion permeation simulations, as evidenced by recent studies. ^41,42^ These factors suggest that the actual energy barrier for Na^+^ permeation could be lower than the one-Na^+^-two-Ca^2+^ PMF estimation, while the actual selectivity ratio might be higher than calculated from the bi-cation permeation simulations. The discrepancies of selectivity ratios from ion permeation and PMF calculations were also observed in previous studies on potassium and sodium channels. ^26,28,36,52^

## 3 Discussion

Fully understanding the ion permeation and selectivity mechanisms of ion channels requires a dynamic picture at the atomic level, which can often be provided by all-atom MD simulations. However, one of the major challenges for calcium channel simulation is the overestimated interactions between Ca^2+^ and the channel. This is particularly true for narrow channels with high selectivity such as Ca_V_1, for which the default CHARMM36m force field would significantly overestimate the Ca^2+^ affinity and underestimate the conductance. To address this, we employed the ECC method with a scaling factor of 0.87, and reproduced the key electrophysiological properties of Ca_V_1, including the conductance (Fig. 1), the mutation effects (Fig. 4), and the valence selectivity (Fig. 5). Notably, the exact scaling factor varies for different studies. ^53,54^ According to the high-frequency dielectric constant of organic materials (ε_el_=2), the scaling factor should be 0.7, ^43^ which yields reasonable results in potassium channels. ^44,45^ Conversely, simulations in prokaryotic sodium channels have demonstrated reasonable conductance without scaling charges. ^29,52^ Accurately modeling electrostatic interactions is critical for capturing the ion-protein interactions within the SF, and the effective dielectric constant remains a key parameter to determine. In the present study, we determined this parameter to be 0.87 for Ca^2+^ by fitting the experimental Ca^2+^ conductance of Ca_V_1. With this parameter, the simulation results were found to be consistent with experimental measurements that were not used for the fitting, including the conductance sequence of cations, the Ca^2+^ ion binding sites, mutation effects, and valence selectivity. These can be viewed as good validations of the parameter.

In our simulations of Ca_V_1, all glutamate and aspartate residues within the SF were modeled in their deprotonated forms. Recent studies on prokaryotic Na_V_ channels have demonstrated that ion selectivity is closely linked to the protonation state of the SF residues. ^55^ In Ca_V_1, previous research has revealed that at physiological extracellular Ca^2+^ concentrations, which are approximately 2 mM, pH can modulate the channel conductance. ^56^ However, when the Ca^2+^ concentration is sufficiently high, the impact of pH becomes minimal. ^56^ Furthermore, the presence of Ca^2+^ has been shown to induce deprotonation of the SF residues. ^57^ Therefore, for our investigation of Ca^2+^ ion permeation and selectivity under a higher Ca^2+^ concentration, we assumed deprotonated states for all acidic residues.

One of the key questions for ion channel studies is the specific ion-binding sites. In previous studies, researchers proposed that there are two ions in Ca_V_, but with only one high-affinity site, the EEEE locus. ^3^ Our MD simulation results support the two-ion configuration within the SF, while further unveils a more nuanced binding landscape. We identified four distinct binding sites (S1U, S1L, S2U, and S2L) within the SF for the two bound Ca^2+^ ions (Fig. 2A, B). Our findings clarify that, while two Ca^2+^ ions are indeed bound simultaneously within the SF, they can dynamically translocate among the four binding sites. This collective translocation is a novel insight that is pivotal for a comprehensive understanding of the Ca^2+^ permeation cycle. Moreover, we observed that the D706 residue around the EEEE locus is deeply involved in the permeation, thus forming an extended EEEED locus for Ca^2+^ binding. Additionally, the V1 site, formed by the D368 residue in our simulations, corresponds to the recently discovered DCS locus. ^15^ This locus serves as a low-affinity binding site in the outer vestibule, facilitating Ca^2+^ entry into the channel.

Regarding the permeation mechanism of Ca_V_, previous studies proposed the two-ion pore model. ^6,7^ Based on our simulation results, we introduce a refined three-ion knock-on mechanism for Ca^2+^ permeation in Ca_V_1 (Fig. 7A). A similar permeation mechanism has been speculated in a recent study based on the -5e charge of the EEEED locus. ^58^ Here we further demonstrate the mechanism in atomistic details and quantify the energetics for ion permeation. Our results show that there are always one or two Ca^2+^ ions coordinating with the EEEED residues in the SF, constituting three key states for ion permeation: state 1, 2U and 2L (Fig. 2D). The PMF calculations indicate that one or two Ca^2+^ ions are not enough for efficient permeation (Fig. S4A, B). Rather, a third Ca^2+^ ion entering the SF reduces the energy barrier to approximately 14 kJ/mol, enabling rapid ion movement (Fig. 3A). The essence of the three-ion knock-on mechanism is the constant presence of two Ca^2+^ ions around the SF, with the third ion facilitating the exit of the first. Although the non-equilibrium ion permeation simulations and equilibrium Metadynamics simulations generated discrepancies regarding the specific occupancies of the three major permeation states, it is reassuring that both support the three-ion knock-on permeation mechanism for Ca^2+^ ions, and the Ca^2+^ conductance and free energy barrier are in good agreement with experimental observations. In addition, our mutant simulations revealed that D706 and E1101 are the most important residues for efficient Ca^2+^ permeation in Ca_V_1 (Fig. 4A). Positioned at the top and bottom of the SF, these residues create enough space for two Ca^2+^ binding and stabilize the two-ion configurations essential for efficient conduction. This finding underscores the importance of precise residue positioning for ion conduction, a factor often overlooked in models lacking atomistic detail. ^18^

**Figure 7:**
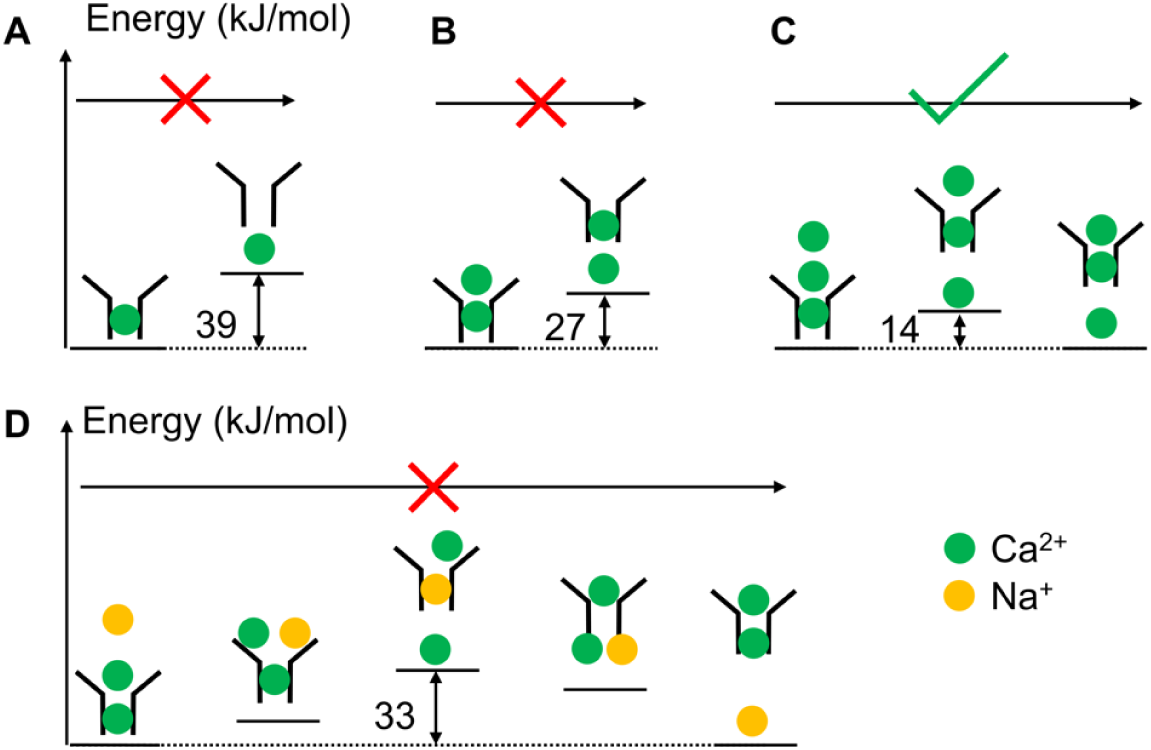
Schematic representation of the Ca^2+^ permeation and selectivity mechanisms in Ca_V_1. (A) A single Ca^2+^ ion is trapped within the selectivity filter of the channel. (B) The presence of two Ca^2+^ ions also fails to promote efficient permeation. (C) The arrival of a third Ca^2+^ ion triggers a knock-on effect that facilitates ion permeation through the selectivity filter. (D) Ca_V_1 exhibits high Ca^2+^ selectivity due to the significantly higher energy barrier for Na^+^ permeation when Ca^2+^ ions are present. The strong electrostatic interactions within the confined SF create an environment that favors the three-ion knock-on permeation of Ca^2+^ over the bypass permeation of Na^+^.

A notable feature of the SF in Ca_V_ is the asymmetry of the EEEE locus, as observed repeatedly in previous studies. Mutagenesis experiments have shown that each of the four glutamate residues contributes differently to Ca^2+^ affinity. ^13^ Additionally, protonation studies revealed an asymmetric binding pattern of proton in the SF. ^59^ This asymmetry has been confirmed in the resolved Ca_V_1 structures, ^21^ with E1406 pointing to the backbone of E364, while the other three glutamates face the pore axis. In our Ca^2+^ permeation simulations, the orientation of E1406 remains stable and rarely coordinates with the permeating Ca^2+^, while the other three glutamates all directly coordinate with Ca^2+^(Fig. 2C). Notably, E1101 also differs from E364 and E705, stably coordinating Ca^2+^ with two carboxylate oxygens (Fig. 2C). In mutant simulations, the roles of these glutamates in Ca^2+^ conduction varied, with E1101 being most important for maintaining the ion conductance (Fig. 4A), which can be explained by the critical position of E1101 for forming the two-ion configurations (Fig. 4C).

While the EEEE locus has long been considered the key residues for Ca^2+^ binding in Ca_V_1, not until recently did researchers identify that the aspartate residue (D706) above E705 is also crucial for Ca^2+^ binding and conduction. ^48,49^ Its role in ion selectivity is minimal though. ^49^ Our simulations demonstrated that the D706 residue coordinates with Ca^2+^ at the S1U and S1L sites (Fig. 2C), alongside E364 and E705. Mutant simulations further confirmed that the D706 is essential for efficient ion permeation (Fig. 4A). Notably, D706 is also highly conserved in the Ca_V_ family. Therefore, we emphasize the significance of the extended EEEED locus in the interactions between Ca^2+^ and the SF of the channel.

Ca^2+^ selectivity is a defining feature of Ca_V_ channels. In physiological conditions, the concentration of Ca^2+^ is far lower than Na^+^ in the extracellular solution, making selective Ca^2+^ permeation a crucial and challenging task for Ca_V_ channels. Previous studies identified the EEEE locus as key to this selectivity,^12,13^ with the mechanism being that once Ca^2+^ binds in the SF, Na^+^ is blocked from permeating. ^6,7^ Our MD simulations reproduced the high selectivity of Ca_V_1 channels (Fig. 5A), confirmed Ca^2+^ blocking, and revealed that the selectivity is from the elevated energy when Na^+^ permeate the channel in the presence of Ca^2+^ (Fig. 6, Fig. 7D). As the transition state showed (Fig. 6C), the high energy barrier for Na^+^ permeation in the presence of Ca^2+^ within the SF results from the inadequate repulsion for pushing the Ca^2+^ to translocate, failing in utilizing the three-ion knock-on mechanism adopted by Ca^2+^ ions. Instead, the permeating Na^+^ has to take a bypass permeation strategy, to translocate by bypassing the bound Ca^2+^ ions with the SF. Even so, the energy barrier is still much higher than the three-Ca^2+^ knock-on permeation, resulting in the high Ca^2+^ selectivity over Na^+^ ions. As for the key residues for Ca^2+^ selectivity, the EEEE locus forms the core single-file binding site for Ca^2+^ and prevents Na^+^ permeation (Fig. 2A). Among these, E1101 appears to be the most important for valence selectivity due to its strong coordination with Ca^2+^ (Fig. 2C) and ability to stabilize two Ca^2+^ within the SF (Fig. 4C), consistent with previous mutagenesis data. ^13^

Similar to Ca_V_, potassium channels also exhibit high ion selectivity, but the structural features and selectivity mechanisms are quite different. Compared with potassium channels, Ca_V_ channels have a much shorter selectivity filter, making bypass events more likely to occur. Our results revealed that in Ca_V_1, Ca^2+^ ions shed 4-5 water molecules in the SF (Fig. S3A), whereas K^+^ ions are fully dehydrated in potassium channels. ^26,27^ The coordination patterns also differ: in potassium channels, K^+^ ions coordinate with the carbonyl oxygens of the backbone, while in Ca_V_1, Ca^2+^ ions coordinate primarily with the carboxylate oxygens of acidic residues, with minimal coordination with carbonyl oxygens (Fig. S3A). These structural distinctions underlie the different selectivity mechanisms. Potassium channels select K^+^ over Na^+^ by sensing different dehydration free energies and ion sizes using carbonyl oxygens, while Ca_V_ channels select Ca^2+^ over Na^+^ by sensing different ion charges using carboxylate oxygens. Despite these differences, both channels share a key similarity: they are multi-ion, single-file channels that can utilize the multi-ion knock-on permeation mechanism. This is a common feature of highly selective ion channels.

There are also less selective Ca^2+^ ion channels, such as the ryanodine receptors (RyRs) and some TRPV channels. For wide-open channels such as RyRs, the selectivity mechanism can be explained by the charge/space competition model. ^17,40^ In RyR channels, the extended region of the vestibule and filter is surrounded by multiple negatively charged residues, facilitating a higher Ca^2+^ occupancy than Na^+^ due to the strong negative electrostatic potential in this confined space. This can generate a weak Ca^2+^ selectivity, but the larger pore radius significantly diminishes the Ca^2+^ selectivity compared to Ca_V_ channels. ^40^ Interestingly, previous studies showed that the weakly selective TRPV1 and TRPV2 channels have one Ca^2+^ binding site with the SF, while the stronger selective TRPV5 and TRPV6 channels have two binding sites with the SF, indicating a correlation between the number of binding sites and ion selectivity.^41,42^ In addition, ion dehydration is also a key factor influencing ion permeability and selectivity. While Ca^2+^ retains its hydration shell in RyR channels, ^33^ it sheds up to 1.5 and 5 water molecules when permeating TRPV6^41^ and Ca_V_1 channels (Fig. S3A), respectively. Greater dehydration indicates stronger interactions with the SF, which could enhance the ion selectivity as well. However, this stronger dehydration also results in lower conductance in TRPV and Ca_V_ channels, necessitating a multi-ion permeation mechanism for efficient ion permeation. Generally speaking, it appears that calcium channels use acidic residues in their SF and outer vestibule to favor Ca^2+^ permeation over monovalent ions, and the number and strength of binding sites within the SF determine the specific ion selectivity.

Although we are able to reproduce many experimental observations and provide new insights and plausible models, there are still limitations in our MD simulations. One potential issue was that the backbone Cα atoms were restrained during the MD simulations. This approach was used because the ion conductance results were hard to converge with a flexible backbone due to the instability of the open-state channel structure, and we were only focusing on the ion permeation and selectivity of the specific open state. Another potential limitation was that our simulations were based on a chimera model using the SF structure of a closed-state Ca_V_ and the gate structure of an open-state Na_V_, which may generate artifacts. Nonetheless, the assumption that the SF structure is similar between the open and closed states appears to be valid, considering the knowledge gained from structural studies of Na_V_ channels. ^60^ Still, these assumptions require further investigation and experimental validation in the future.

## 4 Methods

### 4.1 Simulation systems

The protein models were built using the cryo-EM structure of Ca_V_1.3 (PDB: 7UHG). ^21^ Only the pore domain of Ca_V_1.3, including residue 260-412, 635-760, 1000-1160, and 1340-1472 were included in the simulation systems, and were treated as four chains in simulations. CHARMM-GUI was used to build the simulation systems. ^61^ The protein was embedded into a POPC (2-oleoyl-1-pamlitoylsn-glyecro-3-phosphocholine) lipid bilayer using OPM web server. ^62^ After membrane insertion, the system was solvated in different ionic conditions including 0.15 M CaCl_2_, 0.15 M BaCl_2_, 0.15 M NaCl, and a mixture of 0.15 M CaCl_2_ and 0.15 M NaCl solutions. The simulation box was around 10×10×12 nm^3^, comprising ∼110,000 atoms. The systems and the simulations were summarized in the supplementary tables (Table S1, Table S2). The residue numbers of human Ca_V_1.3 are adopted and those of the key residues are compared with the generic residue numbering system ^63^ in Table S3.

The MD simulations were performed with the program GROMACS version 2021.2, ^64^ using the CHARMM36m force field ^65^ and CHARMM TIP3P water model. For all the production simulations, the time step was 2 fs. The v-rescale ^66^ algorithm with a time constant of 1 ps was used to maintain the temperature at 310 K, and the Parrinello-Rahman ^67^ algorithm with a time constant of 5 ps was used to maintain the pressure at 1 bar. The Particle-Mesh Ewald method ^68^ was used to calculate long-range electrostatics with a cut-off of 1.2 nm, and the van der Waals interactions were smoothly switched off from 1.0 nm to 1.2 nm. The bonds involving hydrogen were constrained using the LINCS algorithm. ^69^ The MDAnalysis package ^70^ was used to analyze the MD simulation results. PyMol ^71^ was used to render molecular visualizations. The default CHARMM-GUI protocol was used to progressively relax the system, including 5000 steps of energy minimization, 250 ps NVT equilibration, and 1625 ps NPT equilibration.

### 4.2 Modeling of the open-state pore

The open pore model was built by adding restraints to the S6 gate region of Ca_V_1.3 channel based on the open-state structure of Na_V_Eh (PDB: 7X5V). ^46^ The four S6 helices of Na_V_Eh were aligned to Ca_V_1.3 for Cα atoms, with the conserved asparagine residues as reference residues. The residue indices for Na_V_Eh are 124-159, with the conserved N144 residue, and the residue indices for Ca_V_1.3 are 380-412, 727-760, 1125-1160 and 1439-1472 for four chains, with the conserved asparagine residues N399, N745, N1145 and N1457. In the MD simulations, we implemented two sets of restraints targeting the protein’s Cα atoms. The first set of restraints targeted residues from the chain’s initiation to a point 5 residues prior to the S6 helix, using the original Cav1.3 structure as the reference coordinates. These restraints were applied with a force constant of 1000 kJ/mol/nm^2^. The second set of restraints was applied to the residues of the gate region, and the reference coordinates for these restraints were derived from the aligned gate region of the Na_V_Eh structure. We performed three successive 0.5 ns simulations with force constants of 10000, 5000, and 1000 kJ/mol/nm^2^ to gradually release the second set of restraints. For the following production simulations, the force constant for the second set of restraints was 1000 kJ/mol/nm^2^.

### 4.3 Force field modification

We used the ECC method to scale the charge of the simulation system. The scaling was applied to the ions and the charged protein residues. In the CHARMM force field, the terminal atoms of a charged residue side chain typically carry a total charge of either +1 or -1. The charges of these terminal atoms were also scaled by the same scaling factor in this study. For nonbonded parameters of calcium, the σ parameter was modified from 0.24357170954 nm to 0.27 nm. This modification was to fit the first peak of RDF (radial distribution function) for the distance between calcium and water oxygen at 0.242 nm. ^72^ Because NBFIX is an alternative strategy to adjust nonbonded interactions like the ECC method, the NBFIX terms for oxygens with calcium and sodium were removed to avoid overfitting. Different scaling factors from 0.82 to 0.89 were tested to fit the experimental conductance of the channel and the scaling factor of 0.87 was found to be optimal.

### 4.4 Ion permeation simulations

A uniform electric field along the negative z-direction (from the extracellular side to the intracellular side) was applied to drive the ion permeation events. We used different strengths of the electric field in the study, varying from 0.0025 to 0.03 V/nm, corresponding to transmembrane potentials of approximately 25 to 300 mV.

The ion conductance (*g*) was calculated by counting the number of permeation events (*N*) for a trajectory in a certain simulation time (*t*) under a transmembrane potential (*V*) as follows:

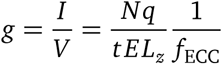

where *q* is the charge of an ion, *L_z_* is the box size in z direction, *f*_ECC_ is the ECC scaling factor. Note that the effective potential sensed by the ions was reduced as charges were scaled in the ECC method. And the charge values *q* used in calculations were integer values without scaling. This is because the ECC scales the charges of the system only to adjust the dielectric constant which does not change the real charge *q*. The errors of conductance were estimated from the standard deviations of the replicates.

The density map of ions was calculated by the “VolMap Tool” in VMD. ^73^ The z-position density and coordination analysis were calculated using MDAnalysis ^70^ package. The origin of the z-position was set to the center of mass (COM) of the Cαatoms of the EEEE locus. The density was calculated as the average number of Ca^2+^ ions in each z-bin. The coordination numbers of oxygens with Ca^2+^ were analyzed with a cutoff of 3 Å for direct coordination. And the averaged coordination numbers were calculated for each z-bin.

### 4.5 Metadynamics simulations

We performed Metadynamics simulations using GROMACS patched with PLUMED ^74^ to quantify the PMF for ion permeation in Ca_V_. We performed simulations for one-Ca^2+^, two-Ca^2+^, three-Ca^2+^ and one-Na^+^-two-Ca^2+^ PMFs. The CV was the z-position of the ion relative to the COM of the Cα atoms of the EEEE locus. The z-position of the ion was restrained by 10000 kJ/mol/nm^2^ walls from -1.3 to 1.3 nm. For the three-Ca^2+^ condition, the restrained region was extended to -2.0∼1.3 nm. The ion motion in the x-y plane was restrained by cylinder restraints with a radius of 1 nm away from the EEEE locus and a force constant of 10000 kJ/mol/nm^2^. For multi-Ca^2+^ sampling, additional walls for Ca^2+^-Ca^2+^ z-distance were applied with a reference distance of 0 and a force constant of 10000 kJ/mol/nm^2^ to avoid crossover of two Ca^2+^ ions in z-direction. Energy Gaussians were deposited every 1 ps with a height of 5 kJ/mol and a width of 0.02 nm. The well-tempered Metadynamics was used with a bias factor of 5. In three-Ca^2+^ sampling, environmental Na^+^ ions were restrained 0.5 nm away from the EEEE locus with a force constant of 10000 kJ/mol/nm^2^ using flat-bottomed restraints. In one-Na^+^-two-Ca^2+^ sampling, other Ca^2+^ ions except the two ions biased by Metadynamics were restrained 1 nm away from the EEEE locus with a force constant of 10000 kJ/mol/nm^2^ using the flat-bottomed restraints. After Metadynamics simulations, the free energy surfaces were obtained using PLUMED ^74^ with a grid spacing of 0.1 Å.

For the 2D- and 3D-PMF results, the MFEP was calculated to project the results onto one dimension, which is the most probable path for ion permeation. The MFEP algorithm is to identify the path connecting two given points with the minimum resistance, which is also called the minimum resistance path. ^75^ The resistance is defined as the integral of the exponential of the energies (exp(*E/k*_B_ *T*)) along the path. ^75^ In principle, we can enumerate all possible paths connecting the two points and find the optimal path, which, however, is computationally impractical for a large network. Here we use fixed points (or energy minima) to coarse-grain the whole state space to a coarse-grained-state (CG-state) network, reducing the computational cost. The algorithm was implemented as follows.

1. Evaluate each grid point to determine which of its neighboring grids has the lowest energy, defining the successor-node list. This list was used to build a whole-space network, where the nodes represent all grid points, and the edges correspond to the successor-node relationships.
2. Identify the fixed points in the whole-space network, which are defined as nodes whose successor node is itself. Each fixed point has an associated connected component in the whole-space network, representing its attraction basin. For each fixed point, we coarse-grained the associated connected component to a CG state.
3. Check the neighboring relationships between CG states by identifying neighboring grids from different connected components. This led to the creation of a CG-state network, where nodes represent different attraction basins and edges represent neighboring relationships.
4. Calculate the MFEP in the whole-space network between every two neighboring CG states, which is defined as the MFEP between the two corresponding fixed points. The shortest path algorithm, implemented in the Python package *networkx*, was used to find the MFEP between the two fixed points by minimizing the resistance.
5. Find the MFEP connecting the two given points. By checking the attraction basins of all CG states, we got the two end CG states whose attraction basins contain the two given points respectively. Applying the shortest path algorithm again to the CG-state network by minimizing the resistance, we obtained the intermediate CG states along the MFEP connecting the two terminal CG states. Connecting the intermediate CG states with MFEPs identified in step 4, we finally obtained the MFEP in the whole-space network.

To reduce computational cost, high-energy regions were excluded from the calculations. After MFEP determination, the mean and error values of PMF were obtained from the average and standard deviation of PMF fluctuation during the last 2 μs of simulations.

## Data Availability

Representative trajectory data with protein and cation coordinates have been deposited in Zenodo and can be previewed with this link. Three supplementary tables (Tables S1–S3) containing the MD simulation summaries and residue numbering systems, and four supplementary figures (Figures S1–S4) illustrating the additional information on the simulation systems, the validation of the simulation system and protocol, the coordination interactions with permeating Ca^2+^ ions, and the PMFs for ion permeation and selectivity (PDF); six movies showing the ion permeation trajectories of Ca^2+^ and Na^+^ ions through Ca_V_1 in mono-cation and bi-cation conditions (mp4).

## Acknowledgements

This work was supported by the Science Fund for Creative Research Groups of the National Natural Science Foundation of China (T2321001). Part of the MD simulations were performed on the Computing Platform of the Center for Life Sciences at Peking University.

## Author Contributions

C.S. and N.Y. conceived the idea. L.X. and C.S. designed the research. L.X. conducted simulations and analyzed data. All the authors participated in the writing of the manuscript.

## Supplementary Information

### 1 Supplementary Tables

**Table S1:**
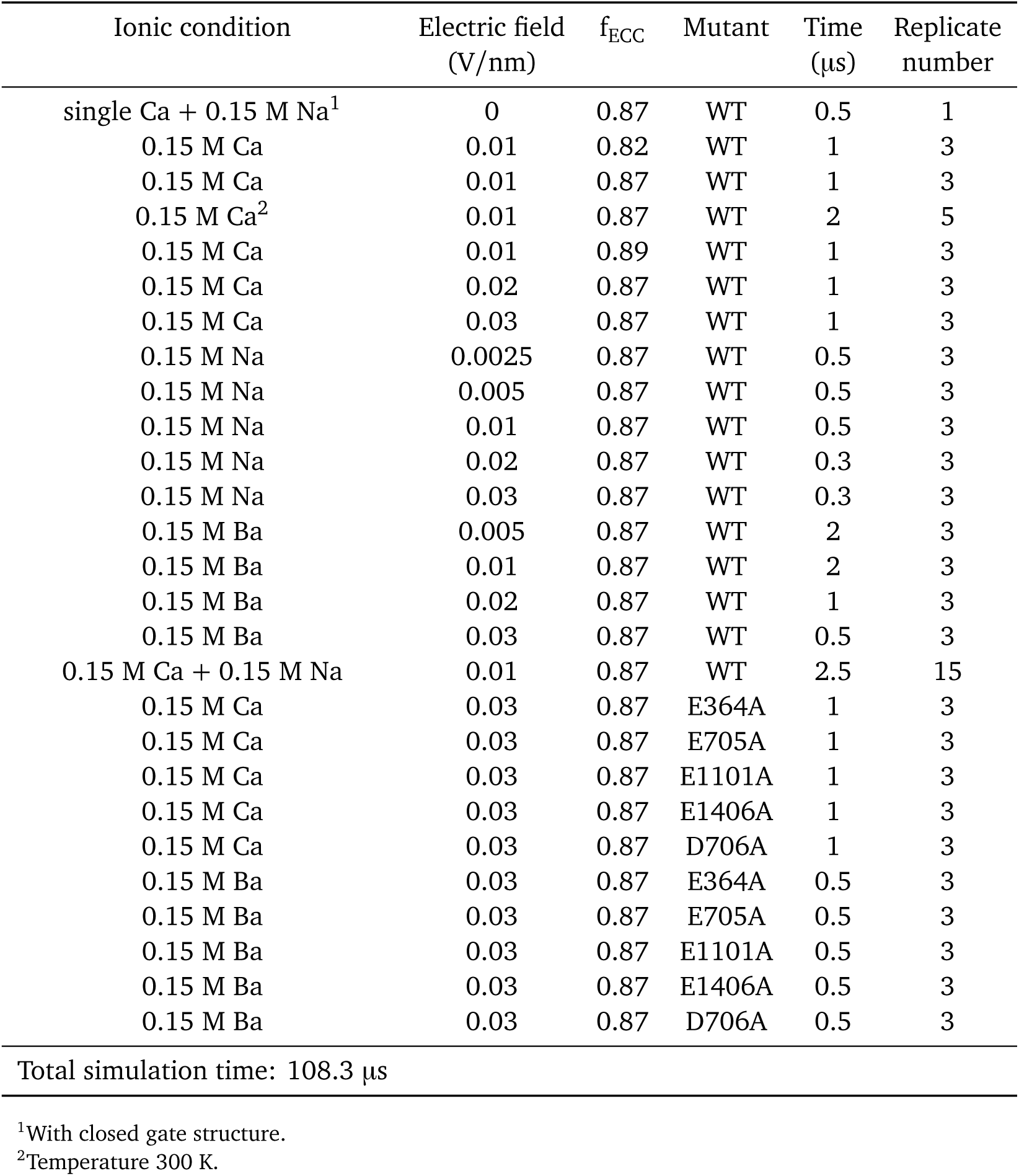
Conventional MD simulations summary.

| Ionic condition | Electric field<br>(V/nm) | $f_{\text{ECC}}$ | Mutant | Time<br>($\mu\text{s}$) | Replicate<br>number |
| --- | --- | --- | --- | --- | --- |
| single Ca + 0.15 M Na<sup>1</sup> | 0 | 0.87 | WT | 0.5 | 1 |
| 0.15 M Ca | 0.01 | 0.82 | WT | 1 | 3 |
| 0.15 M Ca | 0.01 | 0.87 | WT | 1 | 3 |
| 0.15 M Ca<sup>2</sup> | 0.01 | 0.87 | WT | 2 | 5 |
| 0.15 M Ca | 0.01 | 0.89 | WT | 1 | 3 |
| 0.15 M Ca | 0.02 | 0.87 | WT | 1 | 3 |
| 0.15 M Ca | 0.03 | 0.87 | WT | 1 | 3 |
| 0.15 M Na | 0.0025 | 0.87 | WT | 0.5 | 3 |
| 0.15 M Na | 0.005 | 0.87 | WT | 0.5 | 3 |
| 0.15 M Na | 0.01 | 0.87 | WT | 0.5 | 3 |
| 0.15 M Na | 0.02 | 0.87 | WT | 0.3 | 3 |
| 0.15 M Na | 0.03 | 0.87 | WT | 0.3 | 3 |
| 0.15 M Ba | 0.005 | 0.87 | WT | 2 | 3 |
| 0.15 M Ba | 0.01 | 0.87 | WT | 2 | 3 |
| 0.15 M Ba | 0.02 | 0.87 | WT | 1 | 3 |
| 0.15 M Ba | 0.03 | 0.87 | WT | 0.5 | 3 |
| 0.15 M Ca + 0.15 M Na | 0.01 | 0.87 | WT | 2.5 | 15 |
| 0.15 M Ca | 0.03 | 0.87 | E364A | 1 | 3 |
| 0.15 M Ca | 0.03 | 0.87 | E705A | 1 | 3 |
| 0.15 M Ca | 0.03 | 0.87 | E1101A | 1 | 3 |
| 0.15 M Ca | 0.03 | 0.87 | E1406A | 1 | 3 |
| 0.15 M Ca | 0.03 | 0.87 | D706A | 1 | 3 |
| 0.15 M Ba | 0.03 | 0.87 | E364A | 0.5 | 3 |
| 0.15 M Ba | 0.03 | 0.87 | E705A | 0.5 | 3 |
| 0.15 M Ba | 0.03 | 0.87 | E1101A | 0.5 | 3 |
| 0.15 M Ba | 0.03 | 0.87 | E1406A | 0.5 | 3 |
| 0.15 M Ba | 0.03 | 0.87 | D706A | 0.5 | 3 |
| Total simulation time: 108.3 $\mu\text{s}$ | | | | | |

<sup>1</sup>With closed gate structure.
<sup>2</sup>Temperature 300 K.

**Table S2:** Metadynamics simulations summary.

| Condition | Time ($\mu\text{s}$) | Replicate number | Gate state |
| --- | --- | --- | --- |
| One Ca | 4 | 1 | Open |
| Two Ca | 4 | 1 | Open |
| Two Ca | 4 | 1 | Closed |
| Three Ca | 6 | 1 | Open |
| One Na + two Ca | 6 | 1 | Open |
| Total simulation time: 24 $\mu\text{s}$ | | | |

**Table S3:** Key residues of Ca_V_1.3 in conventional and generic residue numbering system.

| Residue | Ca<sub>v</sub>1.3 residue number | Generic residue numbering system |
| --- | --- | --- |
| E | 364 | P2I,28 |
| E | 705 | P2II,28 |
| E | 1101 | P2III,28 |
| E | 1406 | P2IV,28 |
| D | 706 | P2II,29 |
| D | 368 | P2I,32 |
| N | 399 | S6I,30 |
| N | 745 | S6II,30 |
| N | 1145 | S6III,30 |
| N | 1457 | S6IV,30 |

### 2 Supplementary Figures

**Figure S1:**
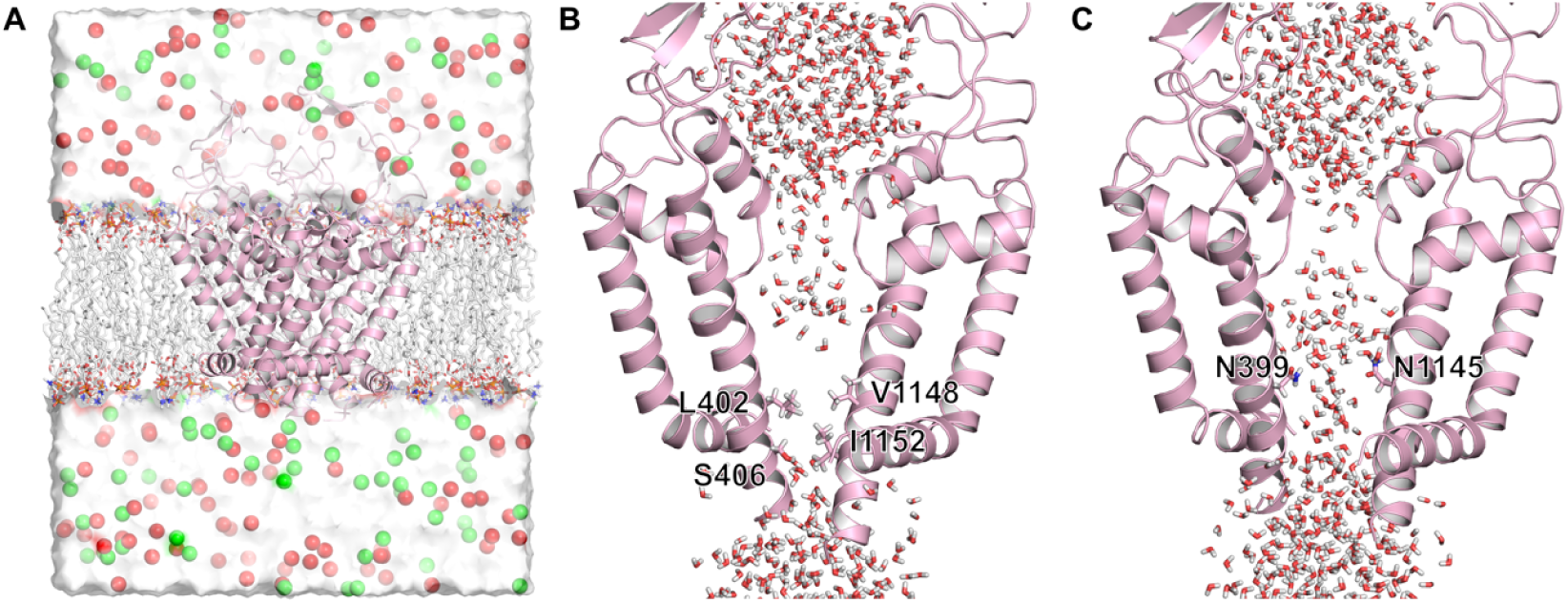
Modeling the open-state structure of Ca_V_1.3 for MD simulations. (A) System overview. The Ca_V_1.3 protein is shown with cartoon. POPC lipids are shown with sticks. Water is shown as white surface. Calcium and chloride atoms are shown in green and red spheres. (B) Side view of the channel with a closed gate. The Ca_V_1.3 protein is shown with cartoon, with repeat II and IV removed for clarity. Side chains of key residues around the intracellular gate are shown as sticks, with residue numbers labeled. Water molecules around the pore axis are shown as sticks as well. (C) Same as (A), but after adding restraints to the S6 gate region according to the open-state Na_V_ structure. The gate region is hydrated after dilation.

**Figure S2:**
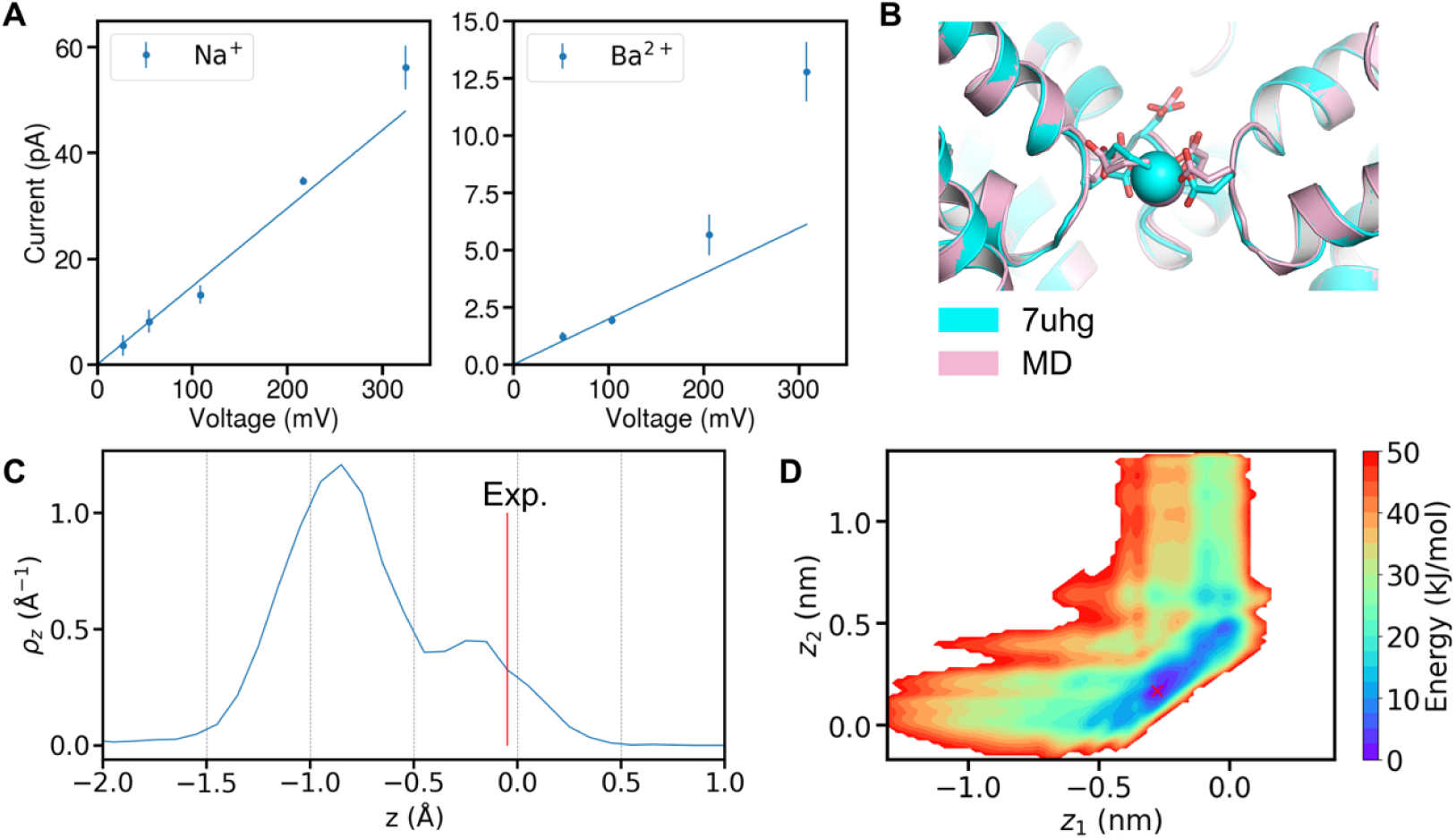
Further validation of the simulation system and protocol by electrophysiological and structural features. (A) The current-voltage relationship for Na^+^ and Ba^2+^. The blue lines represent the results from linear fitting through the origin. (B) The representative SF configuration in one-Ca^2+^ simulations (pink) compared with the experimental structure (cyan) (PDB: 7UHG). The two structures are aligned by C*α* atoms of SF. (C) The z-distribution of Ca^2+^ in MD simulations, with the red line representing the experimental density peak. The reference position is the center of the EEEE locus. (D) Two-Ca^2+^ PMF from Metadynamics simulations, with the red cross indicating the most stable configuration.

**Figure S3:**
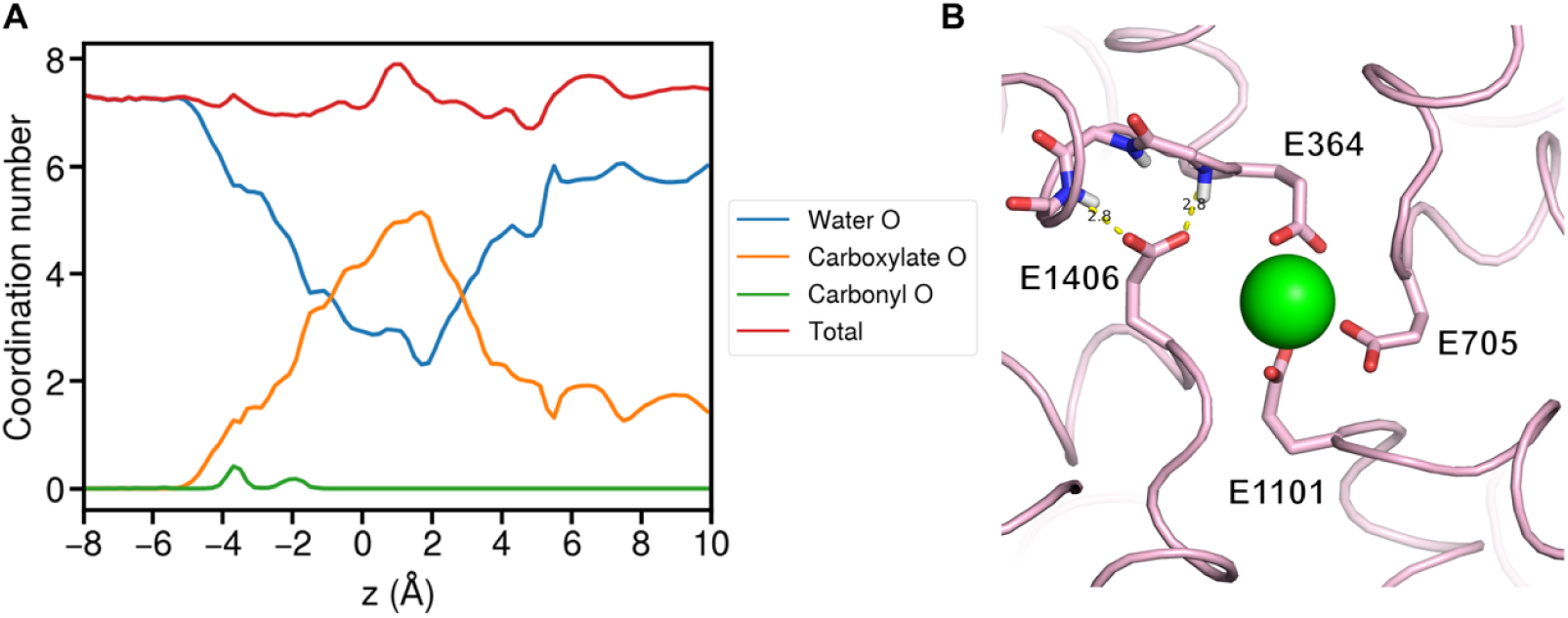
Coordination interactions with permeating Ca^2+^ ions in ion permeation simulations. (A) The coordination number distribution of Ca^2+^ with oxygen atoms along the z-axis. The blue, orange, green and red lines represent the coordination with water oxygens, carboxylate oxygens, carbonyl oxygens, and all oxygens, respectively. (B) Top view of the asymmetric coordination of the EEEE locus. The protein structure is shown as ribbons, with the side chains of the EEEE locus shown as sticks. The green sphere represents the bound Ca^2+^. The hydrogen bonds formed between the backbone nitrogens and the E1406 residue are indicated.

**Figure S4:**
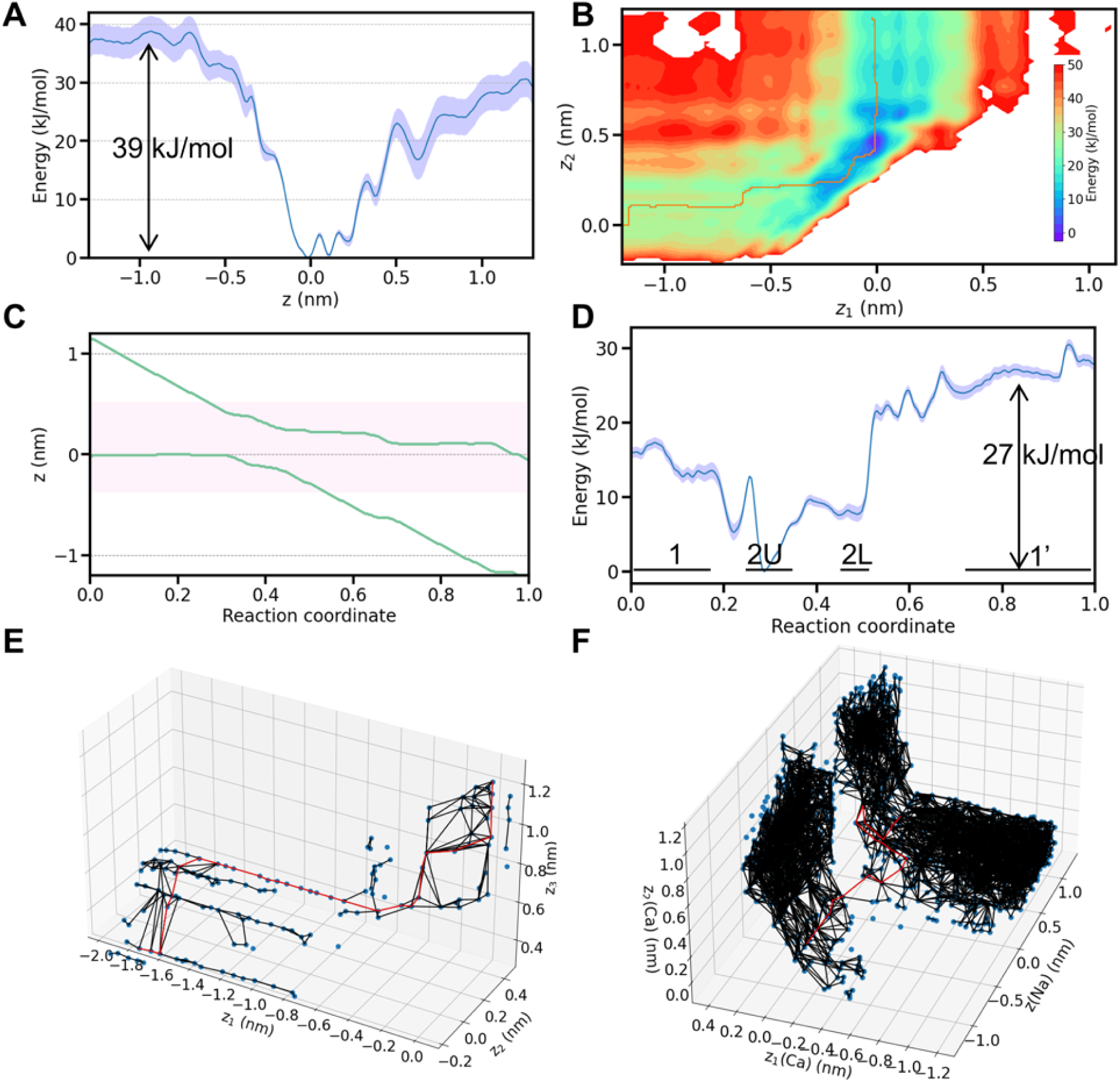
PMF results for ion permeation and selectivity. (A) One-Ca^2+^ PMF profile, with the blue shaded area representing error bars. The arrow indicates the energy barrier for Ca^2+^ permeation. (B) Two-Ca^2+^ PMF results, with the minimum free energy path (MFEP) indicated as the orange line. (C) The coordinated motion of two Ca^2+^ ions along the z-axis of the MFEP. The pink shaded area indicates the SF region. (D) The free energy profile along the MFEP of the two-Ca^2+^ PMF, with labeled states according to positions in (C) and the blue shaded area representing error bars. The arrow indicates the energy barrier for Ca^2+^ permeation. (E) Three-Ca^2+^ PMF results with an energy cutoff of 16 kJ/mol. The local energy minima are shown as blue nodes, with the connecting paths shown as black edges. The nodes and edges with a higher energy than the cutoff are not shown. The red line represents the MFEP. (F) Similar to (E), but for one-Na^+^-two-Ca^2+^ PMF with a cutoff of 36 kJ/mol.

### 3 Supplementary Movies

**Movie S1**

A Ca^2+^ permeation trajectory through Ca_V_1 in pure-Ca^2+^ condition.

**Movie S2**

A Ca^2+^ permeation trajectory through Ca_V_1 in bi-cation condition.

**Movie S3**

A Na^+^ permeation trajectory through Ca_V_1 in pure-Na^+^ condition.

**Movie S4**

The first Na^+^ permeation trajectory through Ca_V_1 in bi-cation condition.

**Movie S5**

The second Na^+^ permeation trajectory through Ca_V_1 in bi-cation condition.

**Movie S6**

The third Na^+^ permeation trajectory through Ca_V_1 in bi-cation condition, in which two Na^+^ translocated together.

